# An encoding of genome content for machine learning

**DOI:** 10.1101/524280

**Authors:** A. Viehweger, S. Krautwurst, D. H. Parks, B. König, M. Marz

## Abstract

An ever-growing number of metagenomes can be used for biomining and the study of microbial functions. The use of learning algorithms in this context has been hindered, because they often need input in the form of low-dimensional, dense vectors of numbers. We propose such a representation for genomes called `nanotext` that scales to very large data sets.

The underlying model is learned from a corpus of nearly 150 thousand genomes spanning 750 million protein domains. We treat the protein domains in a genome like words in a document, assuming that protein domains in a similar context have similar “meaning”. This meaning can be distributed by a neural net over a vector of numbers.

The resulting vectors efficiently encode function, preserve known phylogeny, capture subtle functional relationships and are robust against genome incompleteness. The “functional” distance between two vectors complements nucleotide-based distance, so that genomes can be identified as similar even though their nucleotide identity is low. `nanotext` can thus encode (meta)genomes for direct use in downstream machine learning tasks. We show this by predicting plausible culture media for metagenome assembled genomes (MAGs) from the *Tara Oceans Expedition* using their genome content only. `nanotext` is freely released under a BSD licence (https://github.com/phiweger/nanotext).

## Introduction

The content of its genome constrains the functions an organism can perform. Yet, the definition of function and its representation remain elusive^1^. A genome can be abstracted as a sequence of protein domains, or by analogy as a document containing words. When words are basic units of “meaning”, protein domains are basic units of function. Embedding protein domains in a vector space captures even subtle aspects of this function. The embedding can extended to entire genomes. This results in topic vectors which *distribute* a genome’s functions across latent features^2^. The topic of a document might be “half sport, half politics”. By analogy, the topic of a genome represents its metabolic constraints. Topic or genome vectors encode evolutionary and ecological relationships. They can be used as direct input to learning algorithms. This enables large scale metagenomic applications such as biomining and genotype-phenotype mapping.

In metagenomics, the bottleneck of discovery has shifted from data generation to analysis. Many current sequencing efforts generate huge amounts of data. It is not uncommon to reconstruct thousands of unknown genomes in a single study^3–5^. This wealth of data holds tremendous potential. From the discovery of new enzymes and metabolites to models that predict disease. Related patterns can be detected by powerful learning algorithms such as neural nets^6^. Learning is most effective when the signal in the data is stable across samples. For example, the functions a microbial community encodes tend to be more stable than the taxa it contains^7,8^. To “fit” metagenome-derived functions into learning algorithms, two questions need to be answered: How is function defined? And how is it represented?

We define function as a sequence of protein domains. Most proteins have two or more domains and the nature of their interactions determines a protein’s function(s)^9^. While amino acids build proteins, protein domains represent basic units of “meaning”: They evolve independently^10^ and their structure is often more conserved than their sequence^11^.

Several methods exist to represent function. Most commonly, they use protein domain counts or a gene presence-absence matrix^12–14^. But none of those methods retains context information. Not only can context help to identify protein domains^15^; in microbial genomes, context is vital. Genes in gene clusters are often co-located in *polycistronic* open reading frames (ORFs)^16,17^. Adjacent ORFs can even transcribe as a single mRNA in a form of co-regulation^18^. Most protein domain representations have another disadvantage. They are high-dimensional and sparse, which hinders downstream learning. For example, the *one-hot-encoding* of a Pfam protein domain^19^ has about 17 thousand dimensions, with all elements zero except one.

A representation that preserves context and uses dense vectors are *word embeddings*^20,21^. They assign words with shared context in a document to similar regions in vector space. This assumes that shared context implies shared “meaning”. Word embeddings capture precise syntactic and semanic relationships such as synonyms^22^. Word embeddings assign each distinct word (*vocabulary*) in a large text collection (*corpus*) a vector of real numbers. The most popular training algorithm is `Word2Vec`^22^. It extends to entire documents to create topic vectors^2,23^. Word vectors are a popular input for learning algorithms because they ease training. Here, the embedding acts as a pre-trained language model. The learning algorithm can leverage this knowledge and focus on the task of interest. Embeddings have been trained on biological objects such as genes^24^ and proteins^25^. Yet, because they model the primary sequence, abundant sequence variation will act as noise and hinder training.

Our aim was to encode genomes using protein domains and to use them as input for learning algorithms. The encoding should have several properties: It should represent genome content, not external genome labels such as taxonomy. It should preserve known evolutionary and ecological distances between genomes. The representation should not be sensitive to small sequence variations or incomplete genomes. Formally, it should be a low-dimensional, dense vector of real numbers. Finally, downstream learning tasks using the representation should predict with high accuracy.

## Results

### A topic model of microbial genomes

The encoding model needs a large collection of protein domains (*corpus*) to train on. We collected about 750 million `Pfam` domains from 150 thousand genomes in the *Genome Taxonomy Database* (GTDB, release r89)^26^. Ten thousand domains are distinct (*vocabulary*). Their frequency is comparable to English: Several protein domains are abundant and present in all genomes, like the ABC transporter (PF00005). Other domains are rare and present in only a handful of genomes. The majority of domains follows an approximately log-normal distribution (Fig. S1, A). The corpus was then used as input to the `Word2Vec` algorithm to train an embedding. This weighted, context-sensitive encoding distributes a genome’s content over a vector of numbers. One can then infer new vectors for genomes not seen during training, such as metagenome-assembled genomes (MAGs).

Many hyperparameters influence training. We performed grid search over 96 combinations of parameters (Fig. S1, A and supplementary table 1). Most critical are parameters that adjust the sampling weights of the domains, i.e. the *unigram distribution* (see methods, Eq. 4). Changes in those parameters result in a spectrum of “core” and “accessory” models. Core models focus on domains shared by many genomes. Accessory models focus on domains that are niche- or strain-specific or undersampled.

### Model selection based on ecotype separation

To select between alternative embeddings, many so-called *tasks* exist, on which these models are evaluated. One such task is for example to recognise synonyms^25,27^. We found only one task to translate to genomes – the *semantic odd man out* (SOMO) task^28^. For a set of words (protein domains), we try to identify the one that does not fit the context. For example, “cereal” is odd in the context “zebra, lion, flamingo”. But this task only scores how accurate word-level vectors are. To score document-level vectors, we propose the *ecotype* task: For a given species, the aim is to separate known ecoypes. Ecotypes are subpopulations which are adapted to specific environmental niches. We obtained published ecotype labels for three representative bacterial species (https://github.com/phiweger/nanotext#data). First, the cyanobacterium *Prochlorococcus* (Fig. 1, C) can grow at a range of depths in the oceans’ surface waters^29^. There are two recognized ecotypes that reflect oceanic niche differentiation^30^ – high-light (HL) and low-light (LL). The HL and LL ecotypes can be further refined into subclades^14^ (Fig. 1, D). Second, *Pseudomonas koreensis* (Fig. 1, B) is a group of related species inside the *Pseudomonas fluorescens* complex. Two ecotypes colonise either bulk soil or the rhizosphere^31^. Third, *Vibrio para-haemolyticus* (Fig. 1, A) is a human gastrointestinal pathogen that lives in warm brackish waters. Subpopulations are associated with different geographic locations^32^.

**Figure 1:**
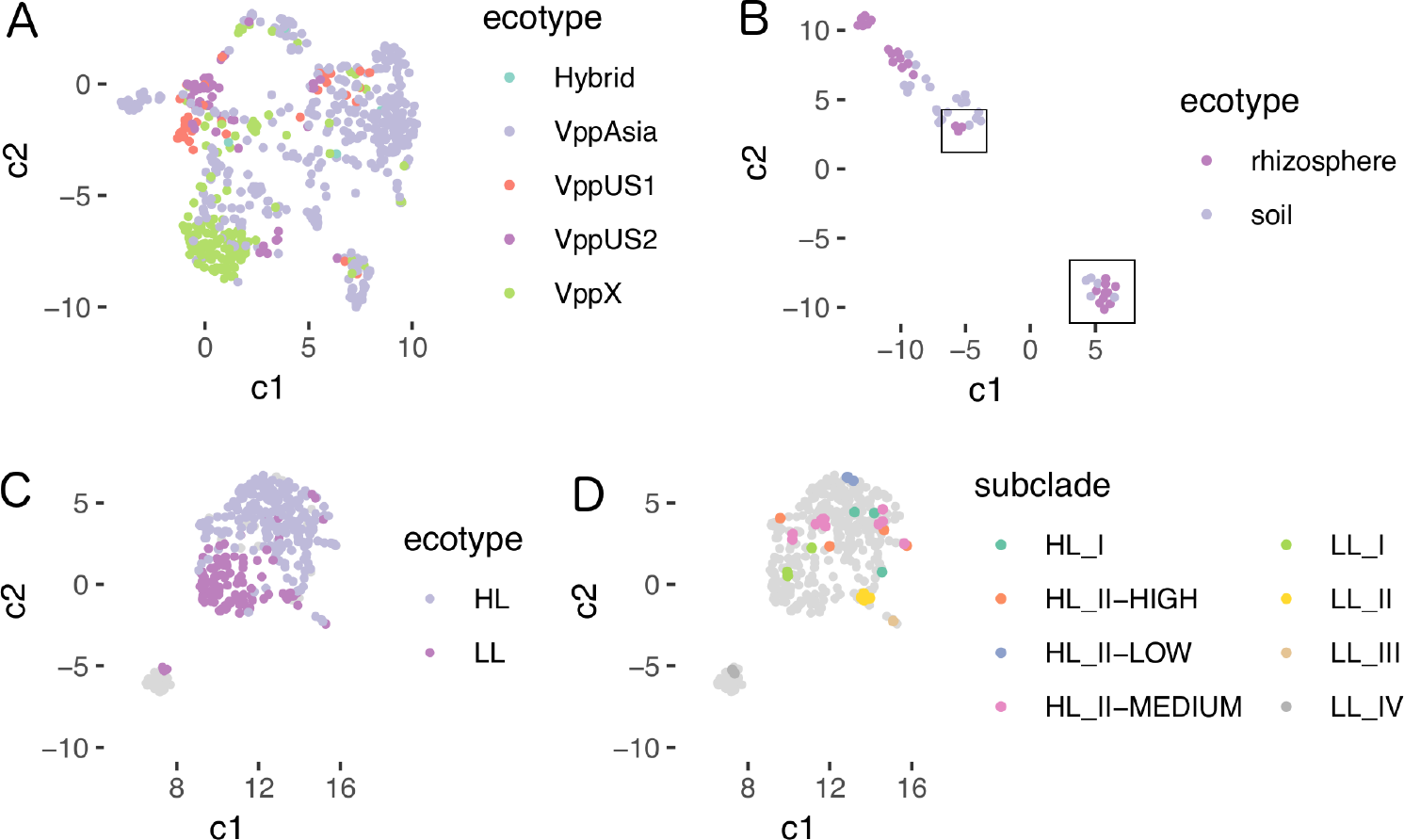
Clear separation of ecotypes using the ensemble model. Vectors were inferred for each genome in the task set and projected into the plane with UMAP^33^. Point colors correspond to ground truth ecotype labels^14,31,32^. **(A)** *Vibrio parahaemolyticus* ecotypes are hardest to separate. They also discriminate best between models. **(B)** *Pseudomonas koreensis* is well separated into its ground truth ecotypes. The original study marked ecotypes by location, not genome content (supplementary figure 3 in *Lopes et al*^31^). This explains why several rhizosphere samples (R15, R23, R24, R45) group with the soil ecotype and vice versa (B2, B3, B7, B11, B35) (boxes). The model distinguishes further clusters within ecotypes, suggesting even deeper niche-differentiation. **(C)** *Prochlorococcus* ecotypes are well separable into high-light (HL) and low-light (LL) ecotypes. **(D)** *Prochlorococcus* ecotypes separate further into subclades.

The SOMO and ecotype tasks helped select three models. Model 93 and 22 correspond to core and accessory models, respectively. Model 45 weights all domains equal. Against a baseline of 16.7% accuracy for random guessing, these three models obtained 84.8, 82.9 and 82.8% accuracy on the SOMO task (Table 1). Generally, models with larger context sizes tend to perform better (Fig. S2). On the ecotype task, the “Vpp” subpopulations of *Vibrio parahaemolyticus* were hardest to separate. This makes them valuable for model selection. Accessory model 22 was best on the ecotype task across all labels (Table 1). Generally, models that give more weight to rare domains outperform core models (S3, supplementary table 2). But they fail when genomes are incomplete. By definition, there are fewer rare domains in a genome than common ones. When the assembly is incomplete, these rare domains are more likely to be missing. This distorts the encoding and results in less accurate genome relationships (see below). Model selection should thus not only rely on task metrics, but on desiderata and on principle, too. As our final model, we averaged the three best-performing models in an *ensemble*. The ensemble model provides clear separations of ecotypes (Fig. 1, A-C). The ensemble model can even distinguish several subclades of *Prochlorococcus* (Figure 1, D). This was not possible in the original study using information on gene presence and absence alone^14^. Our results show that subtle ecological relationships can be inferred from genomic content.

**Table 1:** Accuracy of selected models on the SOMO and ecotypes task. The core model performs best on the SOMO task, while the accessory model excels in ecotype separation.

| model | no. | SOMO | VppAsia | VppUS1 | VppUS2 | VppX |
| --- | --- | --- | --- | --- | --- | --- |
| core | 93 | **84.8** | 86.1 | 19.4 | 56.5 | 77.1 |
| accessory | 22 | 82.9 | **89.9** | **52.8** | **71.7** | **80.7** |
| uniform | 45 | 82.8 | 87 | 38.9 | 65.2 | 79.3 |

### Ecological and evolutionary relationships overlap at higher taxonomic ranks

A fundamental unit of biological diversity is the *species*. Yet, for bacteria and archaea, this concept is subject to ongoing debate^34^. One ecological definition uses shared genome content to estimate how similar two organisms are^35–37^. Shared genome content can also predict phylogeny^38,39^. Whether this is true at the level of species has been questioned^40^.

We quantified the overlap between phylogeny and genome content across all ranks. For this, we compared GTDB and NCBI taxa labels to `nanotext` clusters. The GTDB labels derive from an evolutionary model for 120 ubiquitous single-copy marker proteins^26^. In it, *Parks et al.* extensively reclassified many taxa based on relative evolutionary divergence. `nanotext` is a weighted model of genome content. Its genome vectors form nested clusters which correspond to known taxa. Clusters are more congruent with taxa labels from the GTDB taxonomy, as compared to NCBI (Fig. 2, A and C). Compared with GTDB taxa, coarser clusters correspond to orders (Fig. 2, B), while the most granular clusters correspond to genera (Fig. 2, D and E). At the level of GTDB species, there is little correspondence. `nanotext` clusters contain homogeneous taxa down to the genus level for bacteria and to the family level for archaea (Fig. 3, A). Archaea make up less than 5% of the corpus. Our model has thus too little data to separate archaeal genomes as well as it can separate bacterial ones. It is also too little data to train a separate “archaea-only” model. Our results suggest that metabolic potential and taxonomy are coherent only at higher taxonomic ranks. This was observed recently in an independent study using other methods^40^.

**Figure 2:**
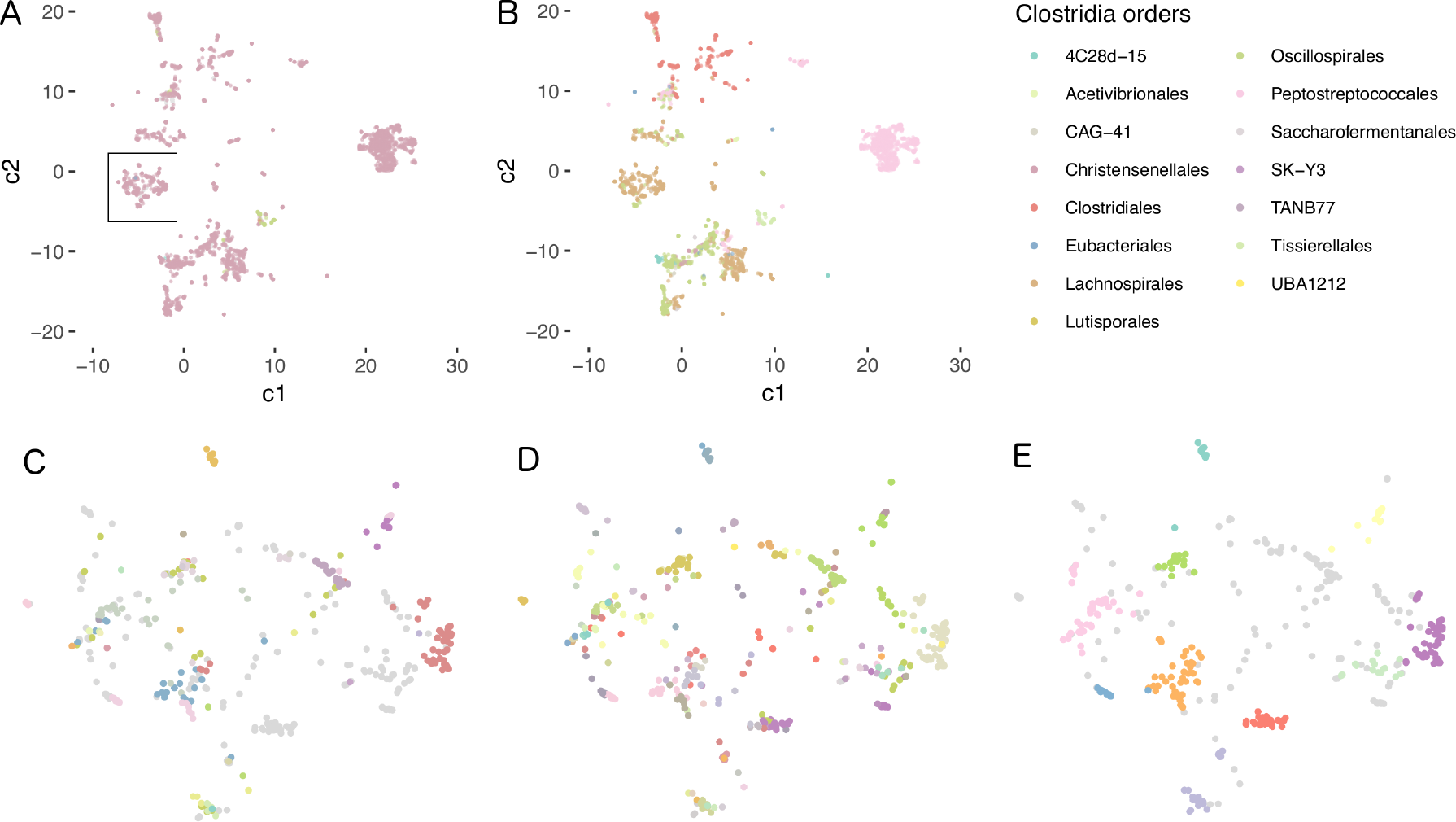
Clustering of genome vectors corresponds to known taxa for ranks down to genus level. Genome vectors from class *Clostridia* were colored by order. *Parks et al.* recently reclassified these orders based on relative evolutionary divergence^26^ (see figure 5 therein). **(A)** The original NCBI taxonomy, where most of class *Clostridia* consists of order *Clostridiales* genomes. **(B)** The reclassified taxonomy released as *Genome Taxonomy Database* (GTDB, r86). It introduces new orders which correspond well to large clusters of genome vectors. **(C)** Detail from A (box); color lables correspond to genera. Many genomes have no assigned NCBI taxonomy (grey). Their clustering matches taxon labels only in part. **(D)** The corresponding GTDB taxonomy is congruent with `nanotext` clusters **(E)**.

**Figure 3:**
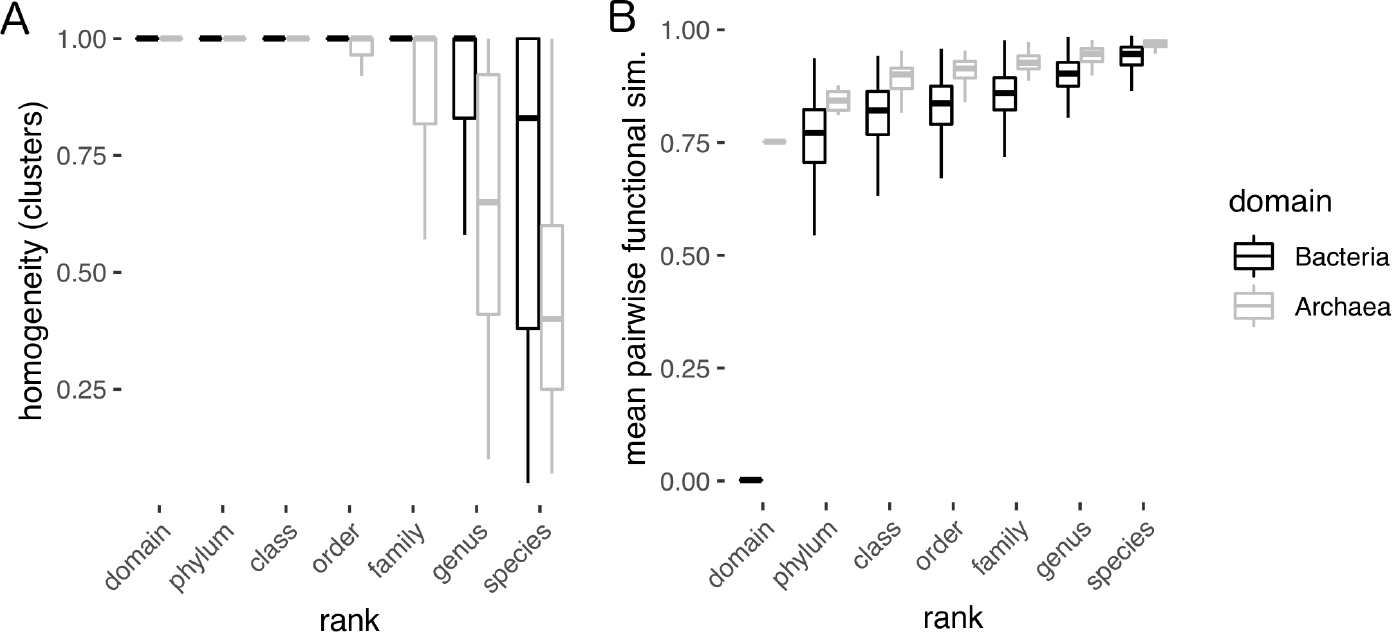
Metabolic potential and taxonomy are coherent at higher taxonomic ranks. **(A)** In the most granular `nanotext` clusters, taxa labels are homogenous at higher ranks. Homogeneity is defined as the fraction of the most abundant taxon within a cluster. At the species level, clusters contain heterogenous labels. This indicates that species fill specific functional guilds. The larger heterogeneity of archaea is an artifact of the low number of archael genomes (< 5%) in the GTDB. Given more data, the model could separate these genomes better. **(B)** The mean pairwise cosine similarity was calculated across all rank labels in the GTDB, i.e. each estimate is calculated “intra-rank” for a given rank label such as “class *Clostridia*”. We downsampled each rank randomly to *n* = 200 genomes. As expected, lower ranks have a high self-similarity compared to higher ranks, approaching zero at the domain level. Again, this effect differs for archaea, due to far fewer genomes in the corpus.

We then “flipped” the analysis to quantify self-similarity within ranks. Here, self-similarity is the average pairwise cosine similarity between pairs of genomes at each rank. As expected, self-similarity increases for lower taxonomic ranks (Fig. 3, B). The average relatedness of any two bacteria approximates null. For archaea, the suboptimal separation of genomes results in a higher self-similarity across all ranks. In summary, `nanotext` clusters are coherent with known taxa at higher ranks. At lower ranks, our model captures ecological features. This is supported by the ability of the model to separate ecotypes (Fig. 1).

### Genome vectors are stable even for highly incomplete genomes

One main use case for genome vectors is to encode MAGs for downstream machine learning. Because most MAGs are incomplete^41^, we assessed the sensitivity of our model to incomplete data. All near-perfect GTDB genomes were selected for the truncation experiment (*n* = 89, 100% complete, 0% contaminated). The GTDB taxa are considered ground truth. Genomes were then truncated *in silico* in 10% increments. Each time, the taxonomy was inferred using sourmash lca^42^ and the `nanotext` core model. `sourmash lca` assigns taxa based on shared k-mers between query and reference genomes. This approach was first popularized by `Kraken`^43^. The `nanotext` core model was chosen because its inference is robust towards incomplete genomes (Fig. S5). It puts little weight on rare domains, which are more likely to be missing from incomplete genomes. For up to 70% missing genome content, both methods returned correct taxa (Fig. 4). `nanotext` alone can correctly assign about 5-10% of taxa. Its accuracy drops at about 70% truncation. `sourmash lca` can classify taxa up to about 90% truncation. Note that both methods are very different. `sourmash lca` can “lock in” on a genome if it contains a set of k-mers that is present in its reference database. `nanotext` is able to learn higher-level features of ranks. This makes it more applicable to searches where few close reference genomes are available. In summary, the `nanotext` genome encoding is robust towards even very incomplete genomes.

**Figure 4:**
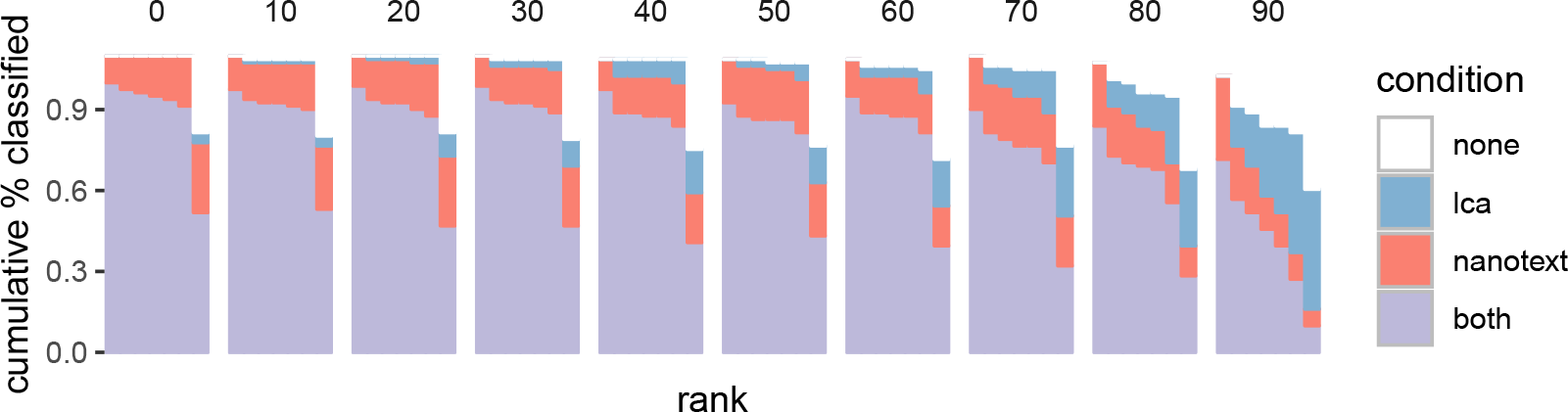
Genome vectors from core model are robust towards incomplete genomes. Near-perfect genomes from the GTDB (*n* = 89) were truncated *in silico* in 10% increments (facets). Each time, both `nanotext` and `sourmash lca` inferred taxa labels across all ranks (x-axis). Either only `nanotext` (red) or `sourmash lca` (blue) were correct, both (violet) or none (white). This resulted in a score for cumulative accuracy (y-axis). Note that about 20% of genomes in the test set had no species label (left most facet, right most bar). `nanotext` is more correct in its assignment of taxa up to about 70% incompleteness. Its accuracy declines thereafter. `sourmash lca` assigns less taxa in general, but does so for up to 90% truncation.

### Functional similarity complements nucleotide similarity

`nanotext` is able to infer relationships for incomplete and distant genomes. This is possible because it complements nucleotide-based distance in two ways. First, `nanotext` uses protein domains for training. These are annotated using *Hidden Markov Models* (HMMs). An HMM in turn distills groups of multiple sequence alignments (MSAs). This aggregates nucleotide variation. Second, `nanotext` embeds the protein domains instead of relying on their names or ID. It places similar domains in similar positions in vector space. This creates the notion of “synonymous” protein domains and aggregates groups of HMMs. `nanotext` quantifies the relationship between any pair of genomes with “functional similarity”. This metric is defined as the cosine similarity between genome vectors. Functional similarity complements nucleotide-based distance metrics like *Jaccard* similarity^44^ and average nucleotide identity (ANI)^45^.

We illustrate this with a use case, where we aim to identify MAGs of an understudied organism with few reference genomes. We search the *Tara Oceans* MAG collection by *Delmont et al.*^4^ for members of the phylum *Gemmatimonadota*. Little is known about this phylum: Most cultured members live in soil^46^. Recently, marine isolates were discovered that photosynthesize^47^ using a non-canonical mechanism^48^ aquired by horizontal gene transfer (HGT)^49^. We first retrieved all genomes labelled *Gemmatimonadota* from the GTDB (*n* = 64). We then overlayed the corresponding vectors with inferred vectors for the *Tara* data (core model, Fig. 5, A). A MAG was returned as putative *Gemmatimonadota*, if it had a cosine similarity *cos* (*θ*) ≥ 0.7 to any GTDB record of the same phylum, a conservative threshold.

**Figure 5:**
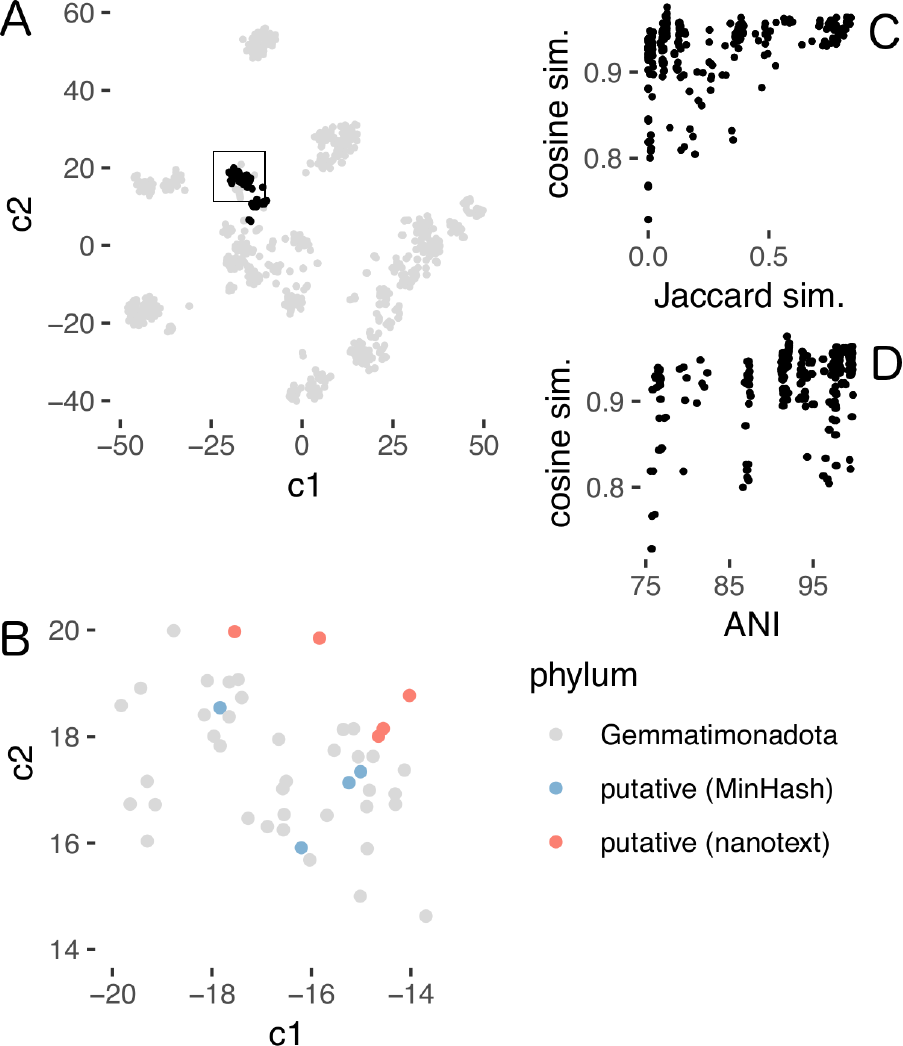
Functional similarity between genomes generalizes nucleotide similarity. **(A)** Members of the understudied phylum *Gemmatimonadota* were indentified from a large *Tara Oceans* MAG collection^4^. We overlayed vectors for all *Gemmatimonadota* in the GTDB (black, *n* = 64) with vectors inferred from *Tara* MAGs (grey, *n* = 957). The GTDB genomes form two distinct clusters. **(B)** Detail of A (black points in box now in grey). Putative *Gemmatimonadota* in the *Tara* collection were identified using functional and nucleotide similarity. `nanotext` and `sourmash` both identified the same five MAGs (red). Additionally, the nearest neighbors identified by `sourmash` based on nucleotide similarity using the `MinHash` algorithm are shown in blue. Note how these are further from the identified MAGs (red) than are the closest GTDB genomes in vector space (grey). `nanotext` can relate these closest genomes with similar metabolic potential even when their nucleotide similarity is low. **(C, D)** Pairwise nucleotide distance between reference and putative *Gemmatimonadota* shows large variance. Yet functional similarity remains stable even past common thresholds of nucleotide-based relatedness (ANI < 0.8, *Jaccard* simiarity < 0.5).

Any similarity search based on exact sequences of nucleotides or amino acids requires close genomes in the reference database. `nanotext` does not share this restriction. Because it relates genomes by content, they can have very different sequence compositions. If their metabolic potential is similar, functional similarity will remain high. We found five genomes that pass our threshold (`TARA_ANE_MAG_00005`, `TARA_RED_MAG_00040`, `TARA_ION_MAG_00042`, `TARA_RED_MAG_00069`, `TARA_RED_MAG_00065` – for additional metadata, see supplementary table 3 in *Delmont et al.*^4^ and http://merenlab.org/data/tara-oceans-mags/). None of these MAGs was assigned a taxon in the original, marker-gene based study^4^. We confirmed these assignments using k-mer based ANI estimation^42^, *in silico* DNA-DNA hybridization^45^ as well as with a phylogenetic approach^26^ (Fig. 5, C and D; supplementary table 3). In summary, functional similarity can relate genomes that have very different nucleotide compositions.

### Accurate prediction of plausible culture media from genome content alone

To test our genome encoding on a machine learning task, we trained a model to predict culture media for MAGs based on their genome content alone. Many unculturable bacteria are thought to be culturable^50^. But it is infeasible to test thousands of medium recipes. Yet, many media share a significant number of ingredients. We therefore trained embeddings of ingredients^51^ for each medium in the catalogue of the *German collection of microorganisms and cell cultures* (DSMZ)^52,53^. To train the genotype-phenotype mapping, we had to link the GTDB to the DSMZ `BacDive` database^53^. We then used a fully-connected neural net to learn this mapping.

Given a genome vector, we were able to predict plausible culture media with high accuracy (Fig. 6, A). The prediction result is a medium vector. Related media can be retrieved via nearest neighbor search. A common-sense baseline is to always predict the most common label of the data set (medium no. 514), which would result in an accuracy of 17%. We classify a prediction as correct, if the target medium is within the top (1, 10) closest media by cosine similarity. This is a common evaluation scheme in multi-class image labelling tasks^6^. On the test set, our model obtains a top-1-accuracy of 63.5% and a top-10-accuracy of 82.5% (Fig. 6, A). On *Tara* MAGs, we obtained a top-1-accuracy of 50% and a top-10-accuracy of 73.2% (Fig. 6, B). The lower accuracy on the Tara data is likely due to genomes without a close representative in the media training data.

**Figure 6:**
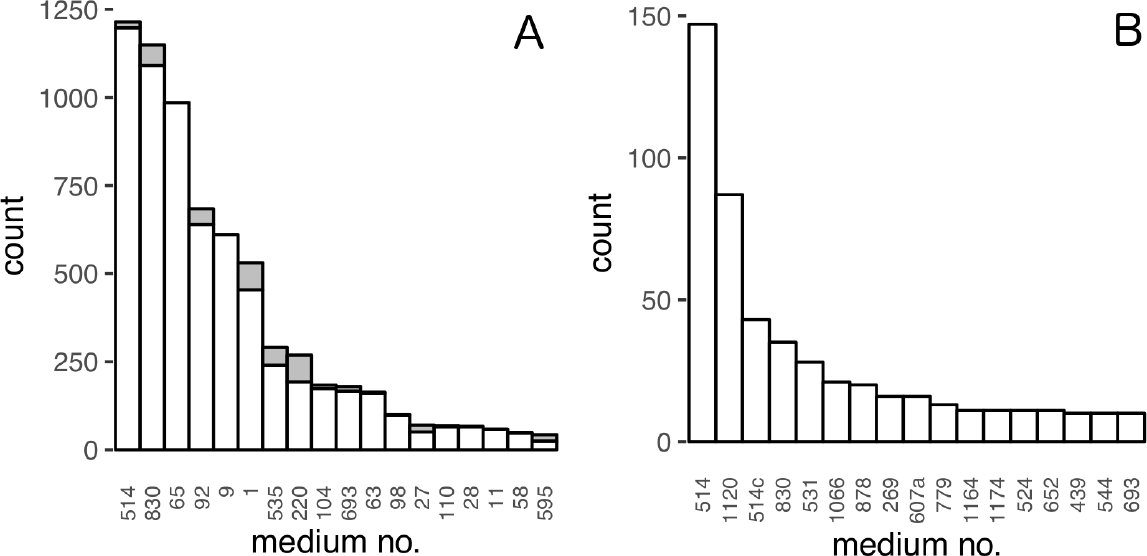
Genome embeddings allow accurate predictions when used as input to learning algorithms. **(A)** Prediction of culture media from genome content. We linked GTDB genomes to DSMZ media at the level of genus. We then trained a fully-connected neural net to learn the genotype-phenotype mapping. Media (x-axis) can be predicted with high accuracy (y-axis, white bars count correct cases, grey bars count wrong cases). A classification was correct, if the the target medium was among the 10 nearest neighbors of the predicted medium vector. Only the 20 most common media in the database are displayed. **(B)** Predicted top media for *Tara* MAGs. The most common media (excluding their variants) in the prediction set are no. 514 (“Bacto Marine Broth”), no. 1120 (“PYSE Medium”) – e.g. used to study *Colwellia maris* isolated from seawater^54^, no. 830 (“R2A Medium”) – developed to study bacteria which inhabit potable water^55^, no. 1066 (“Marinobacter Lutaoensis Medium”), no. 878 (“Thermus 162 Medium”), no. 269 (“Acidiphilium Medium”) and no. 607 (“M13 Verrucomicrobium Medium”) – which includes artificial seawater as an ingredient. All these media correspond to marine isolates. They are plausible starting media for the query MAGs.

A good model should generalise to unseen genome-media pairs. We assessed this on a *Prochlorococcus* MAG (`TARA_ION_MAG_00012`). There were no records for *Prochlorococcus* in `BacDive` at the time of writing. Yet, there exist established culture protocols such as “Artificial based AMP1 Medium”^56^. We labelled the AMP1 ingredients with the same protocol used for the training data^52^. We then inferred a medium vector by summing over the ingredient vectors. Of the 10 nearest media in our embedding space, medium no. 737 (“Defined Propionate Minima Medium”, DPM) has a cosine similarity of 0.979 to the target AMP1. Half of the AMP1 ingredients are found in DPM, including vital trace elements. Other ingredients that are not shared belong to buffers and could be substituted. In summary, genome vectors can be used to train non-trivial learning tasks. We showed this by predicting plausible media for MAGs as a starting point for culture experiments.

## Discussion

In this work we showed how genome embeddings capture many functional aspects of the underlying organism. The main assumption of our approach is that the function of a genome can be abstracted as a sequence of protein domains. Much like words determine the topic of a document, protein domains act as atomic units of “meaning” in a genome. This view of function is very reductive and more comprehensive definitions exist^1^. For example, we do not consider functional RNAs^57^. Yet, our results suggest that `nanotext` performs well on a variety of tasks. This success is likely due to a focus on bacteria and archaea, where most functions are protein-mediated^58^.

We omit other organisms such as eukaryotes and viruses from the corpus. But these can be included without changes to the method. A major bottleneck in corpus expansion is annotation. Currently most approaches are based on *Hidden Markov Models* (HMMs)^59^. These do not scale well to thousands of genomes. It would be interesting to replace protein domain HMMs with homology-based protein clusters. Such clusters could derive from large protein collections such as *UniRef* ^60^ or the *Soil Reference Catalog* (SRC)^61^. A vocabulary compiled from protein clusters would include many proteins of unknown function. Yet, they could still be used in predictive tasks because they are distinct tokens. But the vocabulary size needs restriction. The `nanotext` embedding was trained with a corpus-to-vocabulary ratio of about 10^5^ : 1. This is in line with current corpora in *Natural Language Processing* (NLP) such as `CommonCrawl` (http://commoncrawl.org/). Once a model has been trained, search can scale to billions of vectors^62^.

For training the embeddings we used the `Word2Vec` algorithm. It is a special case of exponential family embeddings^21^. Other embedding methods could be better suited. For example, a *market basket* embedding might be more appropriate to embed media ingredients. The embeddings could be further enriched by subword information^63^. This character-level training on nucleotide sequences would allow inference on *out-of-vocabulary* words. One main limitation of all latent embeddings is that they are not interpretable. Only relative distances within a training run are meaningful. It is thus difficult to reverse engineer our model to identify important domains.

Because `nanotext` encodes genome content, it captures even subtle ecological features. At higher taxonomic ranks, these features are congruent with evolution-based methods^38,39^. At lower ranks, these features complement phylogenetic ones. It has long been recognized that members of a species can be functionally very distinct^8^. For example, *E. coli* can be commensal, enterohemorrhagic or probiotic to the host. There is enormous genomic variation within this species. This diversity is missed in most approaches that focus on marker-genes within the small core genome^64^. `nanotext` encodes the entire genome, an advantage in metagenomics where genomes tend to be incomplete. `nanotext` also uses functional similarity to relate genomes of similar metabolic potential. This complements methods that are based on nucleotide composition. Relationships between genomes can be discovered even though their nucleotide-similarity is below common thresholds. `nanotext` is thus well suited to encode incomplete, distantly related MAGs to either explore or base learning tasks on.

We also showed how genome vectors can be used in as input to machine learning tasks. We illustrated this by predicting plausible culture media for MAGs with good generalization to unseen data. One problem was the lack of full genome sequences in current phenotype databases. We had to create a mapping between the GTDB genome collection^26^ and the DSMZ `BacDive` database^53^. This compromise likely reduces the predictive power of the learned model. Several strain collections have started to whole-genome sequence their inventory. We thus expect a more accurate model in the future. More generally, learning algorithms can become much more efficient when working from embeddings. The algorithms can focus on the learning task and need not learn a “language model” in parallel. Less training data is needed as a consequence. Existing data can be leveraged for machine learning using `nanotext` in two more ways. First, one can use 16S amplicons and “look up” the corresponding functional vector. This approach is similar to `PICRUSt`^65^. Second, for large collections of metagenomes, HMM-based domain annotation becomes a bottleneck. A comprehensive reference database^66^ offers a shortcut. One can then use fast nucleotide-based methods^44^ to search for similar genomes. Once found, their corresponding genome vector can be retrieved by simple lookup.

In conclusion, we trained a genome encoding that captures function in low-dimensional vectors. It solves the “curse of high dimensionality” of previous sparse encodings. Genome vectors are insensitive to missing genome content and nucleotide variation. Pre-trained `nanotext` embeddings can be used for taxonomic classification, biomining and phenotype prediction.

## Supporting information

Accuracy of models on SOMO task

Accuracy of models on ecotype task

Marker gene based taxonomy of target Tara MAGs

## Methods

### Model training

We used all roughly 150 thousand genomes from the *Genome Taxonomy Database* (GTDB, release r89)^26^ for model training. Because the associated taxonomic assignments for release r89 were not yet available at the time of writing, we used the metadata from the previous r86 release. The taxonomy is largely consistent with the expected r89 one (personal communication). Genomes were annotated using `Pfam_scan.pl` (v1.6, https://bit.ly/2CHXlVI) which resulted in a corpus of 750 million domains. Each line in the corpus is the sequence of `Pfam` protein domains on a contig. Strand information is not preserved. We did not filter any protein domains. The vocabulary has 10,879 domains, about 60% of the domains in `Pfam` (v32)^19^.

We obtained word and document vectors using the `Word2Vec` algorithm^22,23^. We trained over a grid of hyperparameter combinations (see below) with a linearly decreasing learning rate (0.025 to 0.0001) over 10 epochs using the *distributed bag of words* (PV-DBOW) training option as implemented in Gensim (v3.4.0, https://radimrehurek.com/gensim/). The result were 100-dimensional vectors for each domain and genome. The similarity of any two vectors can be calculated using cosine similarity (Eq. 1), with a range from −1 (no similarity) to 1 (identical). Cosine distance is defined over the range (0, 2) (Eq. 2). For inference on unseen genomes, we concatenated the annotation results of all contigs. To reach convergence, inference used 1000 epochs. This resulted in stable vector estimates with a pairwise cosine distance less than 0.01 for repeated inferences of the same genome.

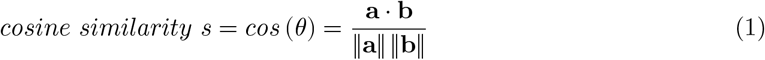

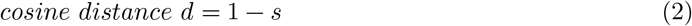

### Hyperparameter optimization

The `Word2Vec` algorithm has many tunable parameters, and they matter^67^. We performed grid search over a range of plausible parameters and trained 96 different models. The most relevant parameters all influence the weight given to individual words based on their frequency (Fig. S1, A). We found that the following parameters influenced training outcome most: the subsampling parameter, the negative sampling exponent, the number of negative samples per frame and context window size (Supplementary table 1).

The *subsampling* parameter *s* determines how likely a word *w*_*i*_ with frequency *z*(*w*_*i*_) in the corpus is kept during training, expressed as a probability *p* (*w*_*i*_) (Eq. 3). Based on this formula, we determined how many of the most frequent domains were affected from subsampling given *s* (Figure S1, B). We set *s* to (3e-5, 1.4e-4) which translates into the most frequent (100, 1000) domains being affected from downsampling.

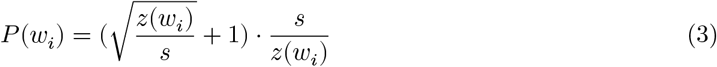

Negative sampling is used during training to teach the neural net which words are not part of the context of a given target word. For efficiency, not all word weights outside the target context are adjusted, but only a small number (between 2 and 20). Negative samples are drawn from a so-called *unigram distribution* (Eq. 4). It scales the count of each domain *f*(*w*_*i*_) by an exponent *v* which was empirically set to 0.75 in the original publications^22,68^. While this parameter is not usually optimized, it can improve model performance on certain problems^67^. Intuitively, *v* provides a handle to focus on less frequent protein domains during training. *v* ≥ 0 attends to protein domains shared by many genomes (“core”) while *v* ≤ 0 focuses on domains particular to only a handful of genomes (“accessory”). For *v* = 0 all protein domains from the “gene pool” are treated equal (Fig. S1, C).

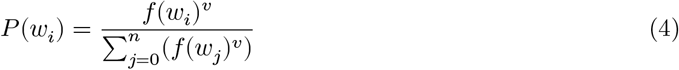

For the remaining parameters the following were used: Window (context) size − (2, 5, 10), number of negative samples (2, 5). The remaining settings were left set to their default values.

### Model postprocessing and nearest neighbor search

After training of word vectors, the average similarity between random pairs of vectors is not zero. Instead, it appoximates a mean vector^69^. We subtracted this mean vector from all model vectors and then unit-normalized them. For fast nearest neighbor search, we used the `Faiss` library (v1.5, https://github.com/facebookresearch/faiss)^62^.

### Task design for model selection

For the SOMO task, we selected one thousand contigs selected at random from the corpus. For each frame of five consecutive domains, one random sample from the vocabulary was added. The mean of the embedding vectors of this set was then calculated. The “odd” domain has the largest cosine distance to this mean. A frame is correct, when the odd domain is the random vocabulary sample. This results in a baseline accuracy of 16.7 % for random guessing. In the ecotype task, each point received the label of its nearest neighbor. If both shared a label, this counted as correct. The average over all points for each ecotype (e.g. soil) resulted in the final score for that ecotype. An accuracy of 1 indicates complete separation of clusters.

### Annotation of Tara genomes

To annotate protein domains for a collection of 957 MAGs^4^ we first identified open reading frames using `Prodigal` (v2.6.3)^70^. We then used `Pfam_scan.pl` wrapping `HMMER` (v3.2.1)^59^ to search against the `Pfam` database.

### Estimation of nucleotide distance and taxonomy

To estimate average nucleotide identity (ANI) between pairs of genomes we used the `MinHash` algorithm^44,71^ as implemented in `sourmash` (v2.0.0a11, https://github.com/dib-lab/sourmash)^42^. The GTDB was used as search index except for the truncation experiments. There, the test genomes were removed before indexing and search. To generate `MinHash` signatures from genomes, we chose a sketch size of `--scaled` 1000 and a k-mer size of 31. For taxonomy inference we used the `sourmash lca` subcommand. We validated all distance estimates using *in silico* DNA-DNA hybridization as implemented in `FastANI` (v1.1)^45^.

### Clustering and dimension reduction

To visualize clusters of genome vectors in high-dimensional space, we projected them into the plane using either the UMAP algorithm^33^ implemented in `umap-learn` (v0.3.7, https://github.com/lmcinnes/umap) for large vector sets or t-SNE^72^ as implemented in `scikit-learn` (v0.20.0)^73^, both with default settings. For clustering we first reduced the vectors to 10 dimensions using UMAP. We then performed clustering using HDBSCAN^74^ as implemented in `hdbscan` (v0.8.19, https://github.com/scikit-learn-contrib/hdbscan) using default parameters and selecting clusters using the conservative `leaf` method.

### Training of media embeddings

To quantify media similarity, we created a media embedding. The current media collection of the DSMZ lists over 1500 media. To reduce the effective number of media, we treated a medium recipe as a sequence of ingredients. We could then use `Word2Vec` to create a latent representation of ingredients^51^. The DSMZ media are not easily parsable and contain many non-unique ingredient tags such as “beef extract” and the synonymous “meat extract”. We therefore used preprocessed data from the `KOMODO` database of known media^52^. To download all 3,637 recipes, we used a custom crawling script (`scrape_komodo.py`). Note that some current additions to the DSMZ media list do not figure in the `KOMODO` database. From each recipe we extracted a list of ingredients^52^. We excluded water (`SEED-cpd00001###`) and agar (`SEED-cpd13334###`) because these ingredients are non-informative. We trained with a window size of 5 and a learning rate as described above over 100 epochs using negative sampling of 15 words per window. To make sure that pairs of media ingredients could occur in the same window, we augmented the data set by shuffling each ingredient list 100 times. The result was a 10-dimensional vector for each media ingredient. To represent a medium, we summed across its ingredient vectors. The similarity of any two DSMZ media could then be compared using cosine similarity. For example, the closest media to medium no. 1 are medium no. 306 (0.99) and no. 617 (0.99), one adding *yeast extract* and the other *NaCl* to medium no. 1; an ID-based representation would treat these media as distict, although they are near identical. Indeed, medium no. 617 and 953 have identical ingredients, which is reflected by a cosine similarity of 1. Any predicted medium can suggest *n* similar media via nearest neighbor search. We visualized the media vector space using t-SNE (Figure S6). Media vectors cluster and enable learning algorithms to discriminate between media classes.

### Linking GTDB genome assemblies to BacDive culture media

To predict a medium from a genome we needed to create a training set that matches the two. The DSMZ BacDive database holds taxonomic and phenotypic information including culture media for currently over 60 thousand strains^53^. However, these strains do not directly correspond to genomes in the GTDB collection^26^. To link these two, we had to pair records using taxonomy at the rank of genera.

### Culture medium prediction

For the medium prediction task, we used a multi-layer, fully connected neural net. We selected the training data as follows: For each genus used to link the two databases, we first sampled records from `BacDive` at the genus level. Because this data is highly skewed towards medically relevant genera such as *Mycobacterium*, we randomly selected a maximum of 100 records per genus. As target **y**, we used the vector of the most common medium for each genus. For the same genus we then randomly sampled a genome vector from `nanotext` (**x**).

We repeated this process 10 times. Data augmentation is a common practice when training neural nets. It enables the training of more complex models, which then generalize better. Using data augmentation, we can circumvent the need to collect more data by varying the input slightly^6^. We used a total of 73,916 genome-media pairs for training, optimized hyperparameters on a validation set of 3891 (5%) and tested the final model on a holdout set of size 8,646 (10%). The neural net architecture consisted of three fully connected layers with (512, 128, 64) nodes. Before applying the non-linear transformation (rectified linear units, ReLU), we normalized the batches of size 128. After each layer we applied Dropout (0.5, 0.3, 0.1). The output layer has 10 nodes to represent a culture medium vector with 10 latent elements, activated by a linear transformation. We optimized a cosine similarity loss using Adam with a learning rate of 10^−2^over the course of 10 epochs. We did not rescale the variables before training. We implemented the model using the deep learning library Keras (https://keras.io).

### Code availability

All relevant resources to reproduce the major results in this article have been deposited in a dedicated `nanotext` repository (https://github.com/phiweger/nanotext). This includes source code for the `nanotext` library and command line tool, preprocessing and training workflows as well as an introductory tutorial. The trained models, corpus, taxonomy metadata and test data were deposited on the *Open Science Framework* (OSF, https://osf.io/pjf7m/).

## Supplement

**Figure S1:**
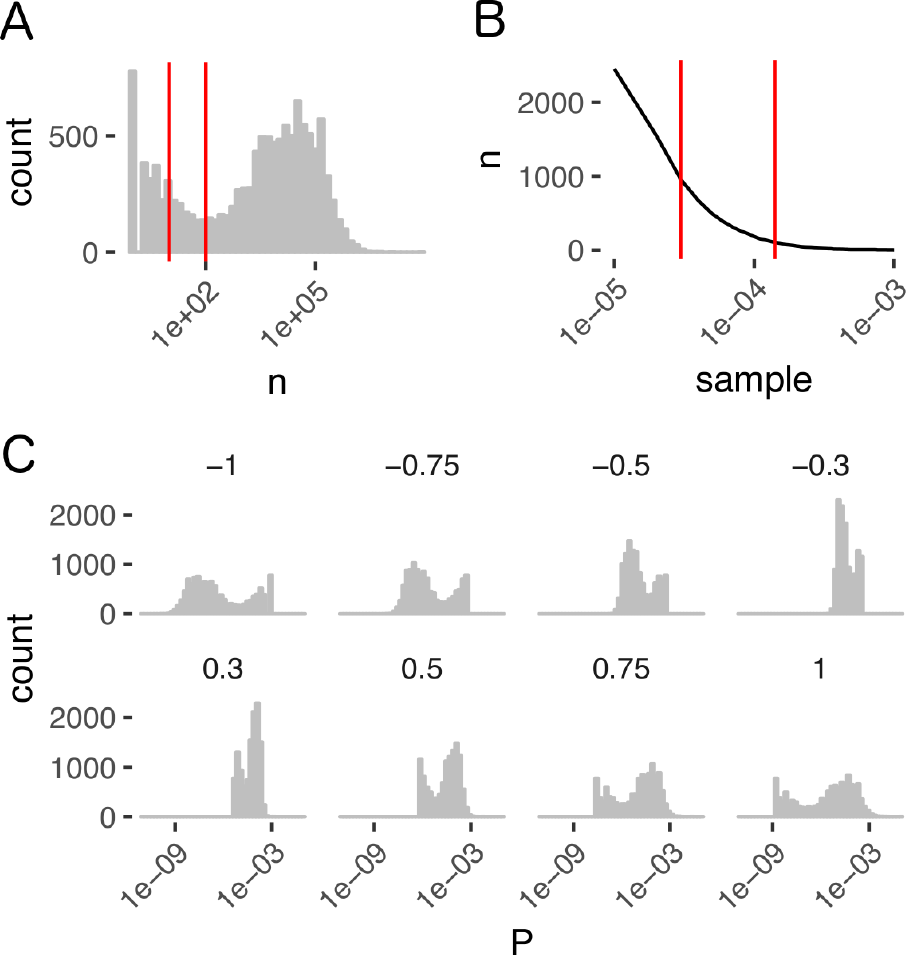
Protein frequency distribution informs hyperparameter selection. **(A)** Count distribution of protein domains. Few domains are present in nearly all genomes while the majority follows a log-normal distribution. There are also many domains present in few copies. Red vertical lines indicate the selected thresholds for minimum count for inclusion in the corpus (left 10, right 100). **(B)** Number if the most frequent domains that are subject to subsampling as a function of different settings for the `Word2Vec` sub-sampling parameter (Eq. 3). Red vertical lines indicate two values chosen for inclusion in the hyperparameter grid search (left 100, right 1000). **(C)** Influence of the negative exponent *v* on the *unigram distribution* (Eq. 4). For *v* = 1 the unigram and frequency distributions are identical. For *v* = −1, they are inverted, i.e. the rarest domains are sampled with the highest probability during training. At *v* = 0, the unigram distribution becomes uniform, i.e. all domains are sampled equal. We chose (−0.3, 0, 0.3 and 0.75 – the canonical default in most applications) for grid search during training.

**Figure S2:**
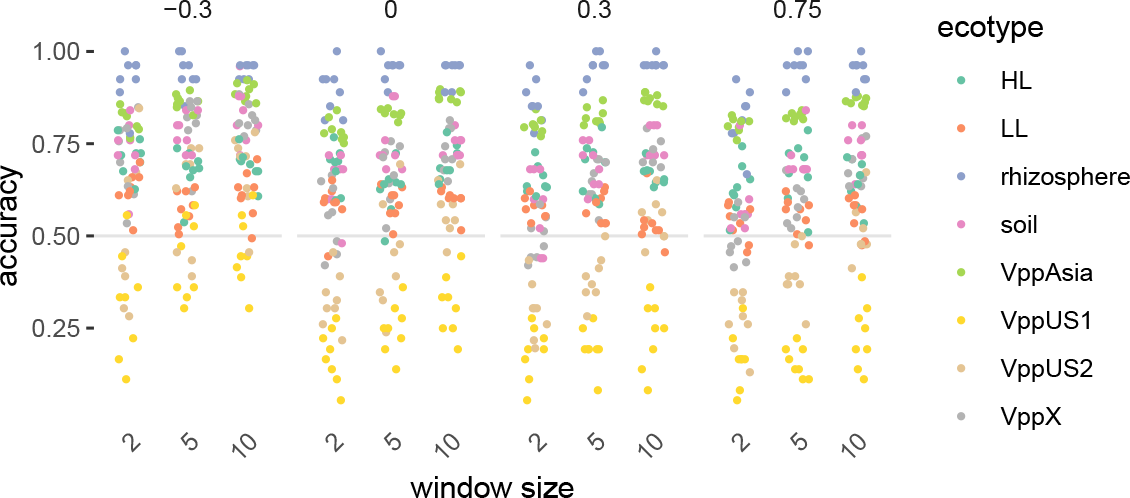
Effect of window size parameter on ecotype task accuracy. Across all negative exponent values for *v* (facets) larger window (context) size performs better for all ecotype labels.

**Figure S3:**
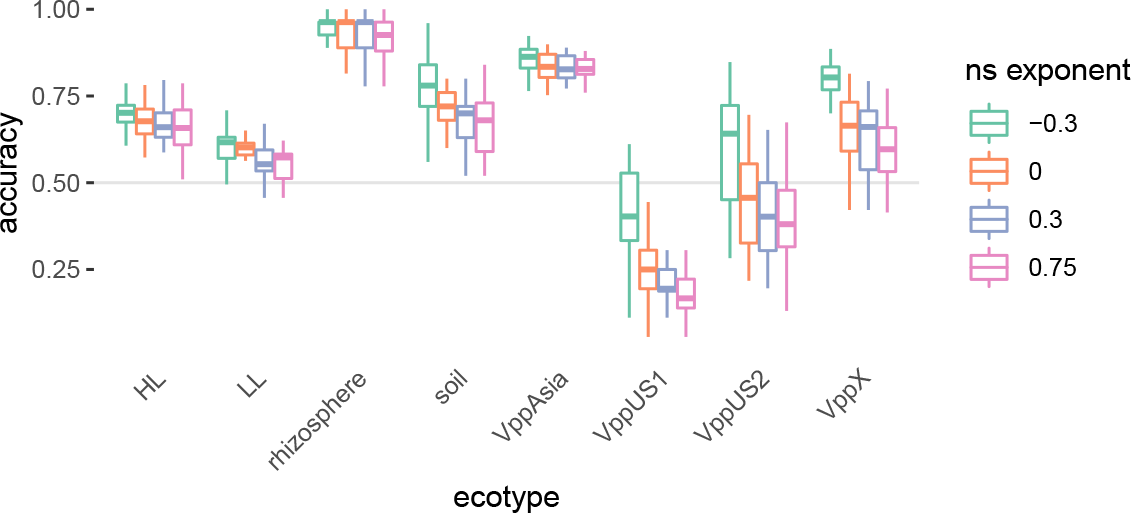
Effect of negative sampling exponent *v* on ecotype task accuracy. Lower exponents lead to more accurate models on the ecotype task across all ecotype labels.

**Figure S4:**
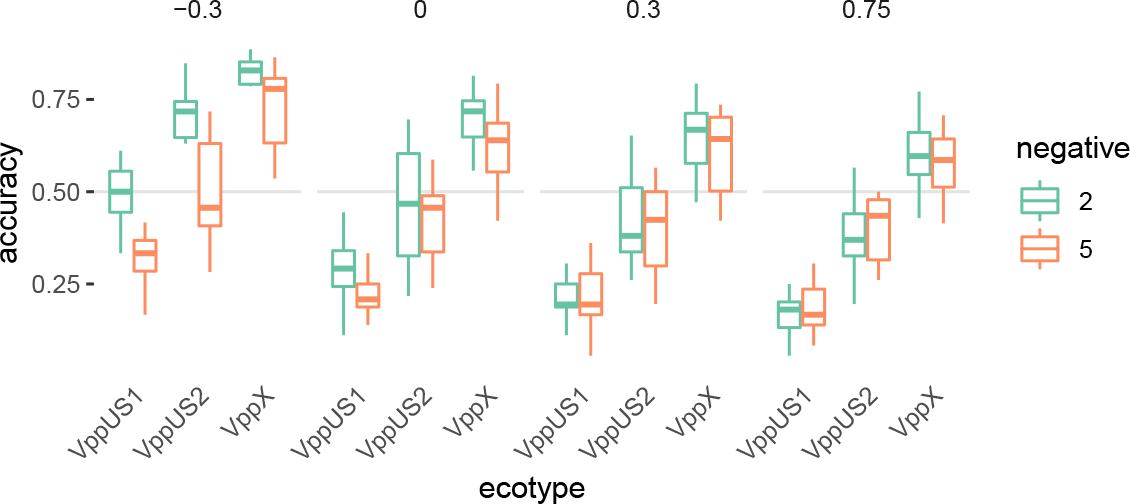
Effect of number of negative samples on ecotype task accuracy. For the hardest to separate and thus most informative ecotypes (x-axis) a smaller count increases accuracy for smaller values of the negative sampling exponent *v* (facets). For *v* ≥ 0.3, the effect is negligible.

**Figure S5:**
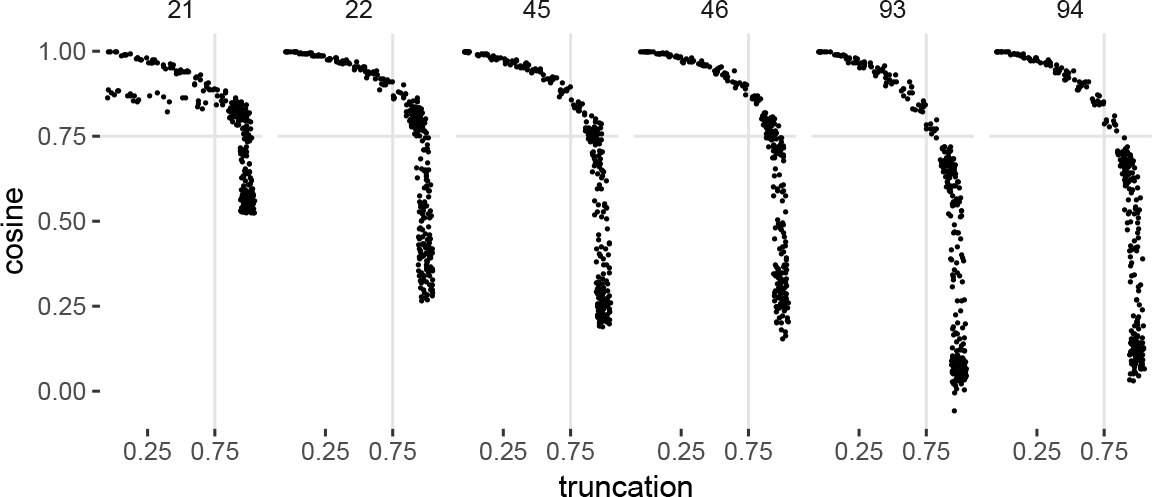
Simulation of genome incompleteness for a single genome (uncharacterized isolate from biogas, unpublished). The genome was truncated in 1% increments (10 repetitions). Each time a genome vector was inferred with a representative model from the hyperparameter grid search (facets). Cosine similarity was calculated against the complete genome. For all training parameters, see supplementary table 1. For accessory models (21, 22), genome vector inference was not stable under truncation. This instability is not observed for models with larger values of the negative sample exponent parameter (45, 46, 93 and 94).

**Figure S6:**
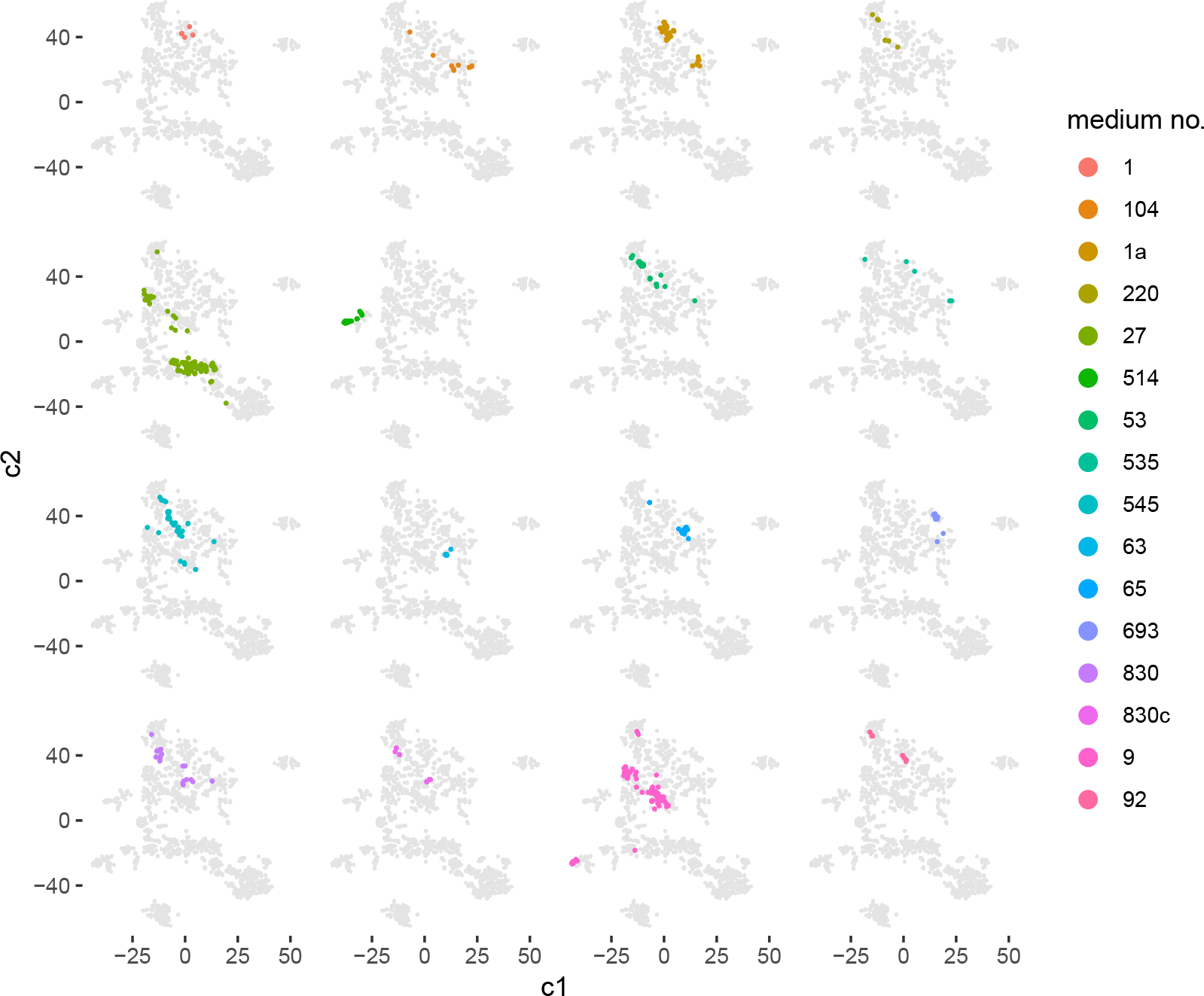
Visualisation of medium embedding space. We used t-SNE to project medium vectors into the plane (grey points). All media with more than 0.95 cosine similarity to any of the top 16 most common DSMZ media were colored. We observe clear clusters of similar media. These clusters can be used by learning algorithms to discriminate media classes. Note how near-identical media such as no. 830 and no. 830c are embedded in near identical vector space.

